# Pan-cancer distribution of cleaved cell-surface Amphiregulin, the target of the GMF-1A3 antibody drug conjugate

**DOI:** 10.1101/2022.03.10.483840

**Authors:** Kristopher A. Lofgren, Nicolette C. Reker, Sreeja Sreekumar, Paraic A. Kenny

**Affiliations:** Kabara Cancer Research Institute, Gundersen Medical Foundation, La Crosse, Wisconsin, USA; Department of Medicine, University of Wisconsin School of Medicine and Public Health, Madison, Wisconsin, USA

**Author notes:** Corresponding Author: Paraic A. Kenny.

## Abstract

Amphiregulin (AREG) is a transmembrane protein which, following TACE/ADAM17-dependent cleavage, releases a soluble Epidermal Growth Factor Receptor ligand domain that promotes proliferation of normal and malignant cells. Expression of Amphiregulin has been described by immunohistochemistry in several tumor types, including lung, prostate, head and neck, gastric, pancreatic and breast cancers but evidence for a functional requirement for Amphiregulin in these malignancies is more limited. We have previously described the development of a monoclonal antibody, GMF-1A3, that selectively recognizes the Amphiregulin epitope that is revealed following cleavage by TACE/ADAM17 and demonstrated that drug conjugates of this antibody have anti-tumor activity in mouse models. By directly evaluating Amphiregulin cleavage, immunohistochemistry on tissue specimens using this antibody can be used to evaluate the extent to which Amphiregulin is being proteolytically processed in cancer, which is a more direct measure of Amphiregulin activity. As a potential companion diagnostic for this antibody-drug conjugate, this immunohistochemistry assay allows identification of tumors with high levels of the cleaved Amphiregulin target. Here we evaluate levels of cleaved Amphiregulin in 370 specimens from 10 tumor types and demonstrate that it is widely expressed in solid tumors and is especially common (more than 50% of cases) in breast, prostate, liver and lung cancer.

## INTRODUCTION

Amphiregulin (AREG) is a transmembrane protein which, following TACE/ADAM17-dependent cleavage [1], releases a soluble EGFR ligand domain which promotes proliferation of normal and malignant cells. This proteolysis event leaves a residual cell-surface transmembrane stalk which is subsequently internalized. We determined the N-terminal sequence of this cell associated Amphiregulin cleavage product [2] and generated antibodies that selectively recognize this epitope in its cleaved but not its intact conformation [3]. The antibodies are internalized by cultured cells in a cleaved Amphiregulin dependent manner [3]. We have developed one of these antibodies into an MMAE-based antibody drug conjugate, GMF-1A3, and demonstrated that it can kill human breast cancer cells *in vitro* and as xenografts in immunocompromised mice [3].

Appropriate selection of patients for targeted cancer treatment typically relies on some kind of companion diagnostic to identify the sub-population of patients whose tumors are most likely to respond. An initial evaluation of our GMF-1A3 antibody as a potential immunohistochemical companion diagnostic was performed in 138 breast cancer specimens. We found medium/high immunoreactivity in 70% of cases [3]. While our initial drug development focus has been on Amphiregulin in breast cancer, it is expressed in several other cancer types [4] so the potential utility of therapeutic antibody drug conjugates likely extends to other malignancies. While the expression of Amphiregulin in these other tissues has been described, the extent to which cleaved Amphiregulin is present at sufficient abundance to potentially represent a viable target for the GMF-1A3-MMAE antibody drug conjugate is unknown. To address this issue, we have evaluated the levels of cleaved Amphiregulin in tissue microarrays comprised of 370 specimens from a total of 10 tumor types.

## MATERIALS AND METHODS

### Immunohistochemistry

Two multiple organ carcinoma tissue microarrays were purchased from BioCoreUSA (Philadelphia, PA, USA). The slides were deparaffinized in xylene and rehydrated by serial incubations in graded ethanol and then in water in a Histo-Tek^®^ SL Slide Stainer (Sakura Finetek USA, Inc., Torrance, CA, USA). Antigen retrieval was performed in a steamer by boiling slides in a container of citrate buffer (pH 6.0) for 20 min which was then removed for 15 minutes of cooling on the benchtop. Slides were washed in 1x Wash buffer (Dako Agilent, Santa Clara, CA, USA) and endogenous peroxidase was quenched by incubating with Dako Dual Endogenous Enzyme Block for 10 minutes. Slides were washed in 1x Wash buffer, blocked (5% rabbit/10% goat serum in PBS), and immunostained with goat anti-AREG antibody (15 μg/mL; AF262, R&D Systems) or rabbit anti-cleaved AREG 1A3 antibody (10 μg/mL) overnight at 4°C. Slides were washed four times in 1x Wash buffer, followed by incubation for 45 minutes at room temperature in 1:100 dilution of rabbit anti-goat immunoglobulins/HRP or ready-to-use goat antirabbit HRP labelled polymer (Dako). The slides were washed twice in 1x Wash buffer and the color was developed with 3,3-diaminobenzidine tetrahydrochloride (DAB) substrate chromogen system (DAKO). Sections were washed with water and counterstained with hematoxylin, rinsed with water, dehydrated by serial ethanol washes to 100%, cleared, and mounted in Permount (Thermo Fisher Scientific). The staining intensity was assessed semi-quantitatively using a four-point scale (Negative=0, Low=1, Medium=2, High=3) by two investigators working independently on blinded samples. Discordant scores were resolved by joint review.

The tissue array design included 380 cores. In the TMA sections used, 370 cores were evaluable for cleaved Amphiregulin (Figure 1) and 369 were evaluable for both cleaved and total Amphiregulin (Figure 2).

**Figure 1.**
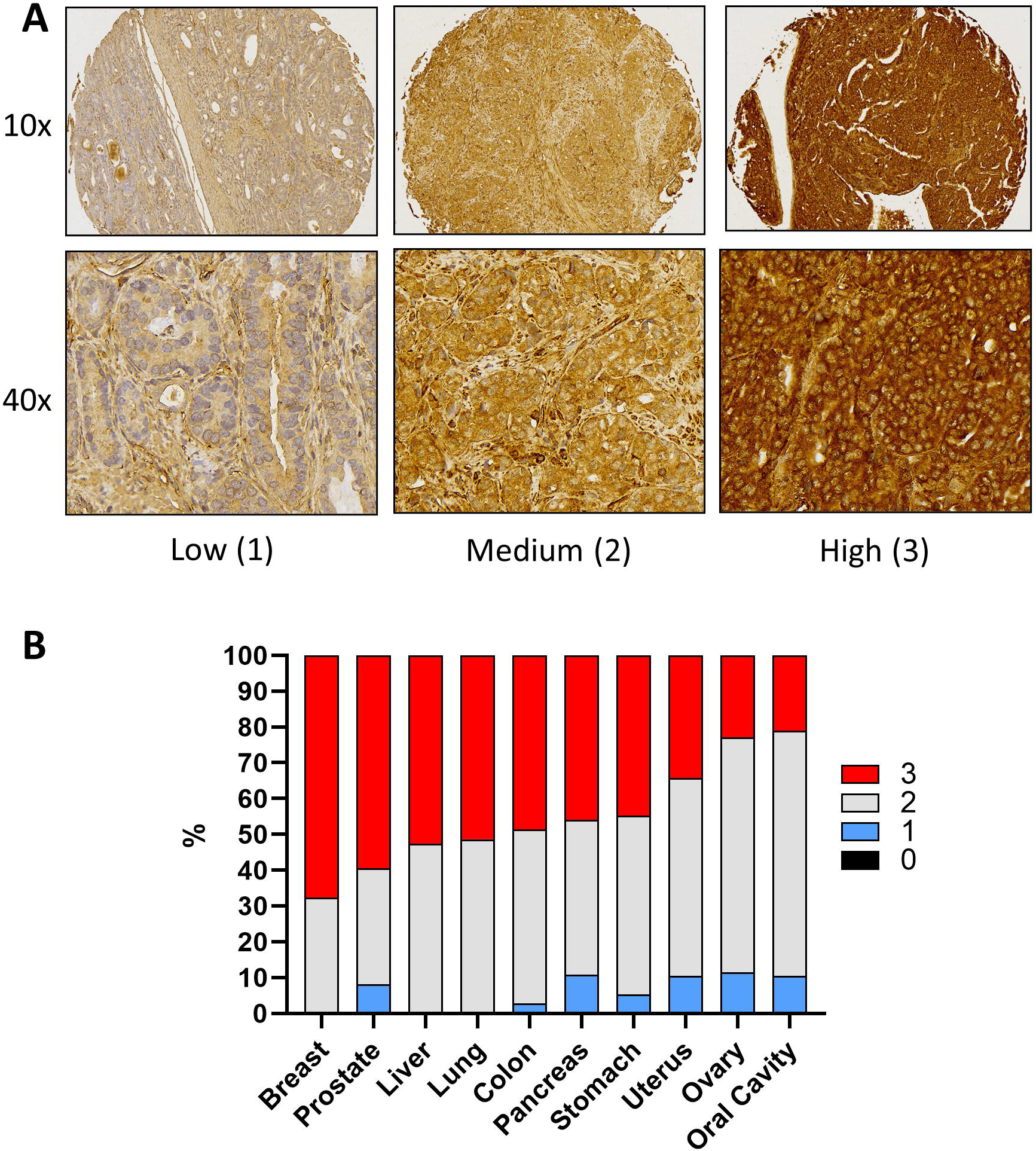
Tissue distribution of cleaved Amphiregulin in solid tumors. (A). Representative examples of immunohistochemical staining for cleaved Amphiregulin at each of the three detected staining intensities. Images from the same cores were captured at 10x and 40x magnification. (B). Quantification of cleaved Amphiregulin intensity scores in 380 tumors from ten different cancer types.

## RESULTS

### Tissue distribution of cleaved Amphiregulin in cancer

To broadly evaluate the levels of cleaved Amphiregulin in cancer, we performed immunohistochemistry on multi-tissue tumor microarrays. The neoplastic compartment of each specimen was scored semi-quantitatively as Negative (0), Low (1), Medium (2) or High (3). No neoplastic cells were negative for 1A3 immunostaining. Representative examples of each staining score are shown in Figure 1A, and data from all tissues examined are summarized in Figure 1B. A circumferential staining pattern was especially evident in the high intensity specimens (Figure 1A, right). At least 50% of breast, prostate, liver and lung tumors stained in the highest score for cleaved Amphiregulin.

### Cross-comparison of levels of total and cleaved Amphiregulin in cancer

The specificity of the GMF-1A3 antibody for cleaved Amphiregulin was previously verified by evaluation of cross-reactivity against peptides representing cleaved and full-length Amphiregulin [3]. If GMF-1A3 antibody immunohistochemistry also accurately reflects cleaved Amphiregulin levels in formalin-fixed tissue then we would anticipate that intensity scores for both GMF-1A3 and total Amphiregulin would tend to be positively correlated and, in particular, that cases with low-to-absent expression of total Amphiregulin should be very unlikely to exhibit robust immunostaining with GMF-1A3. We immunostained the tissue microarray with an antibody against total Amphiregulin and cross-compared the intensity scores for each antibody. Data from all tissues are summarized in Figure 2, and data from individual tissues are shown in Figure 3.

**Figure 2.**
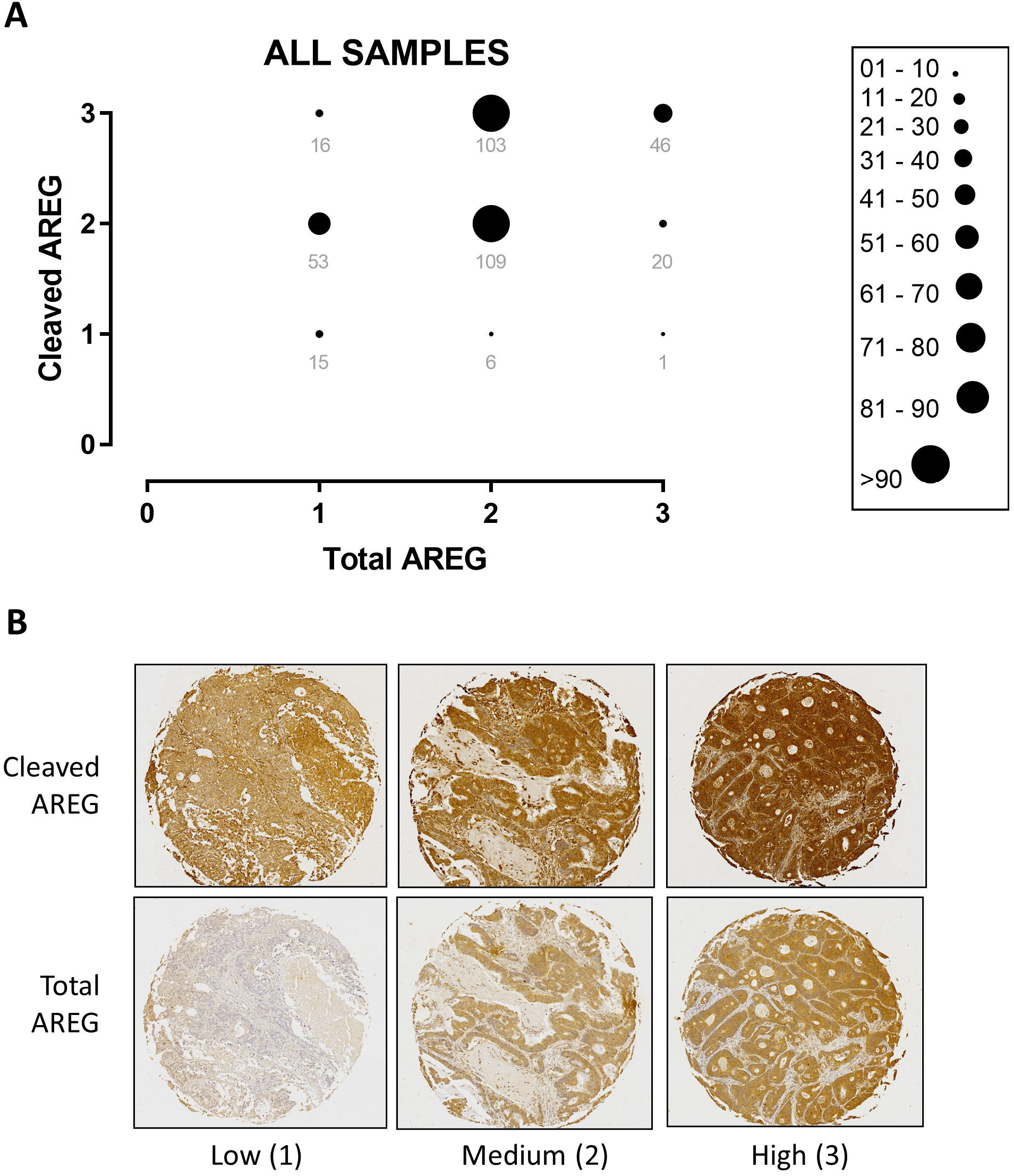
Comparison of levels of cleaved and total Amphiregulin in all 370 tumors. (A). Cross-comparison of intensity scores for both cleaved (Y-axis) and total (X-axis) Amphiregulin in 380 tumors. Data point size is proportional to the number of cases in each pairwise group and the number of cases in each group is indicated. (B). Representative examples of tissue cores in which both total and cleaved Amphiregulin both had low, medium or high immunostaining intensity.

**Figure 3.**
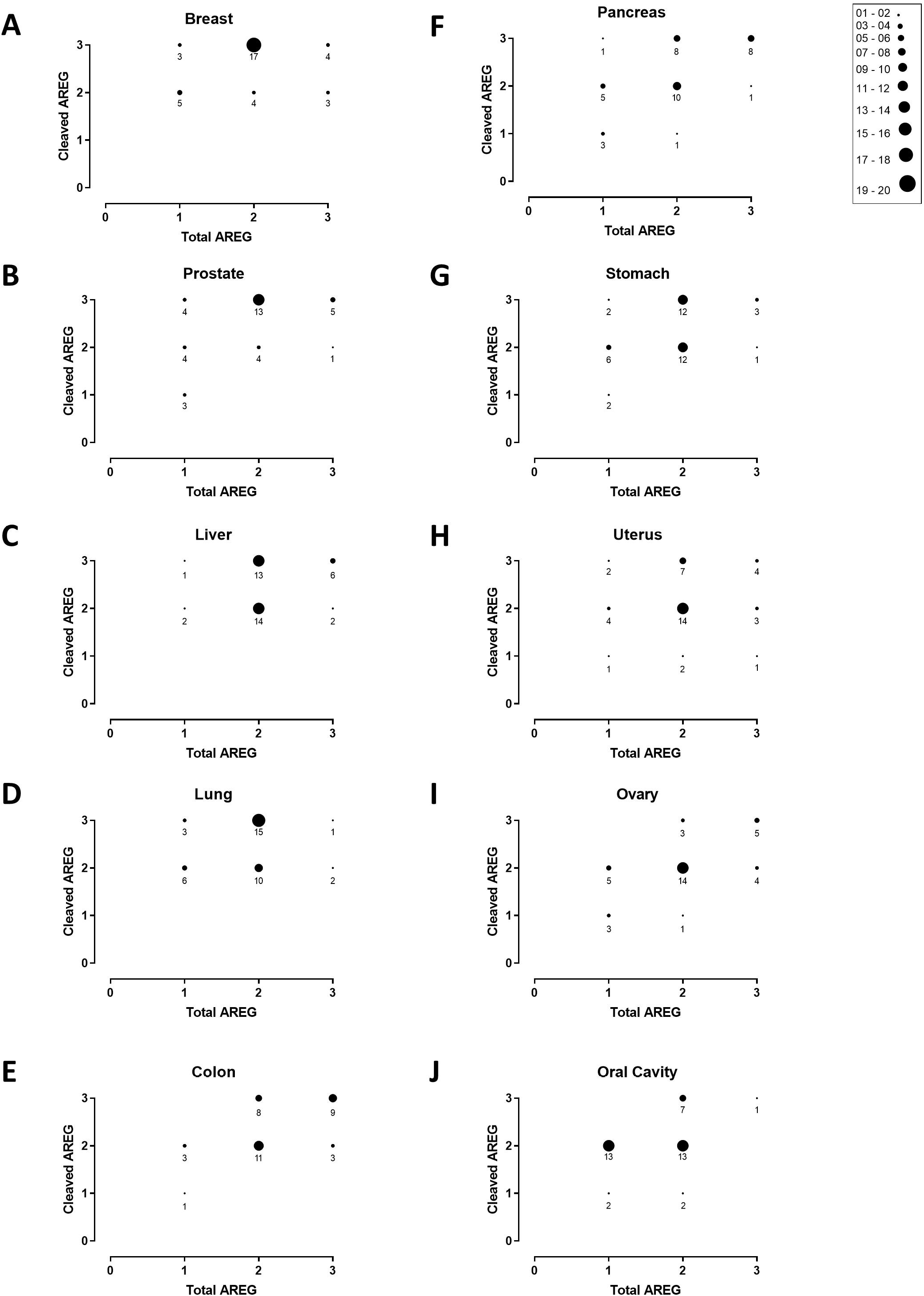
Pairwise comparison of cleaved and total Amphiregulin levels in ten tumor types. (A-J). Cross-comparison of intensity scores for both cleaved (Y-axes) and total (X-axes) Amphiregulin in each of the ten indicated tumor types. Data point size is proportional to the number of cases in each pairwise group and the number of cases in each group is indicated.

As anticipated, there was a generally positive correlation between total Amphiregulin levels and levels of cleaved Amphiregulin. All specimens had at least some Amphiregulin immunoreactivity and only 16/369 specimens had the highest score for cleaved Amphiregulin and the lowest (but not negative) score for total Amphiregulin. We also noted that cleaved Amphiregulin and total Amphiregulin immunoreactivity were found together in the same tissue compartment (e.g. representative example of medium intensity immunostaining in Fig 2B). Where we observed a departure from a linear positive relationship between the pair-wise scores (most prominently in breast, prostate, liver and lung, Figure 3), it was in specimens having the highest intensity score for cleaved Amphiregulin with a medium score for total Amphiregulin, suggesting particularly active levels of Amphiregulin processing in these tumors.

## DISCUSSION

In this study, we extended our prior findings in breast cancer specimens [3] to several additional malignancies, showing that Amphiregulin expression is widespread and, when expressed at moderate to high levels, Amphiregulin cleavage was commonly detected in these specimens. This substantially extends the tissue repertoire where the cleaved Amphiregulin target of the GMF-1A3-MMAE antibody drug conjugate is commonly expressed.

Expression of total Amphiregulin in several of these tissues had previously been described [5–9], but evidence for a functional requirement for Amphiregulin has been limited to a smaller group, including colorectal [10] and breast cancer [6, 11]. Our demonstration here that the expressed Amphiregulin is being actively cleaved in a wide range of cancer types confirms that autocrine Amphiregulin signaling following ADAM17-dependent Amphiregulin cleavage is commonly occurring, raising the possibility that Amphiregulin-dependent EGFR activation may be a more frequent mitogenic signal that has been previously appreciated. Accordingly, in addition to being potentially useful as a companion diagnostic for the GMF-1A3-MMAE, this antibody may facilitate a more complete appraisal of the extent of active autocrine Amphiregulin/EGFR signaling in both normal tissue development and homeostasis as well as cancer initiation and progression.

## ACKNOWLEDGEMENTS

This study was supported by the Department of Defense Breast Cancer Research Program (W81XWH-14-1-0294 to PK), the American Cancer Society (123001-RSG-12-267-01-TBE to PK), and the Gundersen Medical Foundation. KL was supported by the Norman L. Gillette, Jr. Breast Cancer Research Fellowship. PK holds the Dr. Jon and Betty Kabara Endowed Chair of Precision Oncology.

